# Investigating the role of mitochondrial membrane potential in paternal inheritance of mitochondria

**DOI:** 10.1101/2023.10.30.564200

**Authors:** Ariane Pouliot-Drouin, Thierry Niaison, Sophie Breton, Stefano Bettinazzi

**Author notes:** Corresponding author: Stefano Bettinazzi;, UCL Department of Genetics, Evolution & Environment, 99-105 Gower Street, London WC1E 6BT, UK. Statement of authorship:* APD: data analysis, writing-original draft; TN: data collection and analysis; SBr: conceptualisation and supervision; SBe: conceptualisation, methodology, data analysis and supervision. All authors contributed substantially to revisions.

## Abstract

The process of oxidative phosphorylation (OXPHOS) in mitochondria depends on an electrochemical gradient known as the mitochondrial membrane potential (Δ*ψ*m). Reflecting high functionality, elevated Δ*ψ*m usually depicts healthy mitochondria and contribute to organelle selection. This study investigates whether mitochondrial properties linked with bioenergetics, such as Δ*ψ*m, may play a role in paternal inheritance of mitochondria. More specifically, how sperm Δ*ψ*m responds to egg chemoattractants in bivalves characterized by distinct mitochondrial inheritance patterns: strict maternal inheritance (SMI) and doubly uniparental inheritance (DUI), the latter displaying sex-specific transmission of paternal mitochondrial DNA (mtDNA). Sperm Δ*ψ*m was examined in four bivalve species: *Mytilus edulis* and *Ruditapes philippinarum* (DUI), plus *Mercenaria mercenaria* and *Mya arenaria* (SMI). In absence of oocytes, sperm Δ*ψ*m did not vary between the two groups. However, we revealed an increase in Δ*ψ*m following egg detection only in sperm bearing paternally-derived mitochondria (DUI). This suggests, along with bioenergetic changes, that Δ*ψ*m modulation might be a specific property of DUI paternal mitochondria, possibly implicated in their unique ability to be sex-specifically transmitted.

## Introduction

Eukaryotic life depends on mitochondrial respiration to satisfy most of the cellular energy requirements. This is achieved through the process of oxidative phosphorylation (OXPHOS), which relies on an electrochemical gradient generated by proton pumping and substrate oxidation. This intermembrane electric potential (i.e. the mitochondrial membrane potential, Δ*ψ*m) drives ATP production by the ATP synthase complex [1]. Given its fundamental role for energy production, loss of Δ*ψ*m often depicts mitochondrial dysfunction and compromised bioenergetic efficiency, with downstream impact on cellular fitness [2,3] (but see the potential benefits of mild depolarization: [4,5]). Therefore, Δ*ψ*m can be used to discriminate between healthy and dysfunctional mitochondria, and consequently contribute to the selective elimination of underperforming mitochondria by mitophagy to ensure viable organelle transmission [2,6–9].

Strict maternal inheritance (SMI) of mtDNA, a near universal mitochondrial transmission system in animals, might have evolved as a way to prevent transmission of selfish mutations and preserve viable mitochondrial function in the offspring [10,11]. Discarding sperm mitochondria allows the removal of mitochondrial variants which have been potentially exposed to higher oxidative stress [12,13]. Moreover, it promotes homoplasmy (i.e. a single type of mtDNA in a cell or individual), therefore perpetuating efficient mitonuclear interactions [14,15]. Its opposing state, heteroplasmy, is recognized for its general negative effects on embryonic development, fitness, behaviour, cognition, age-related diseases and survival [16–20]. To prevent heteroplasmy, paternal mitochondria and mtDNA must systematically be eliminated by different pre- and post-fertilization mechanisms, such as mtDNA removal during spermatogenesis in *Drosophila melanogaster*, degradation of ubiquitin-tagged paternal mitochondria in bovine and primate embryos or mitophagy in embryos of *Caenorhabditis elegans*, to name a few [19,21–24].

The exception to SMI in animals resides in a little over a hundred bivalve species that possess a very different doubly uniparental inheritance system (DUI), where paternal mitochondria are typically transmitted to sons and populate male gametes, while maternal mitochondria populate all somatic tissues and female gametes [25–28]. In other words, sperm mitochondria escape degradation in male bivalves [29]. The DUI system thus provides a unique opportunity to study the mechanisms involved in paternal mitochondria preservation and transmission to future generations, its possible adaptative value for male functions and even link with sex-determination [30,31].

Previous studies suggested that the genetic divergence between female- and male-derived mtDNAs (F-mtDNA and M-mtDNA, respectively) result in the expression of different mitochondrial phenotypes, underpinning sex-specific bioenergetic adaptations with potential repercussion on fitness [32–35]. The male-specific mitochondrial phenotype in DUI is characterized by a strong limitation exerted at the end of the electron transport chain, resulting in limited respiratory rates [32,33,35], a trait potentially in line with a high membrane potential in male mitochondria. Knowing that mechanisms of mitochondria quality control can exploit Δ*ψ*m to discriminate highly functional organelles [3], a putative role of Δ*ψ*m in driving paternal mitochondria persistence in DUI populations has been suggested [34,36,37]. However, whether Δ*ψ*m contributes to mitochondrial selection and might represent one mechanism underpinning doubly uniparental inheritance of cytoplasmic organelles remains to be demonstrated.

In this study, we took one step further in our understanding of the putative role played by sperm Δ*ψ*m in paternal inheritance of mitochondria. We examined mitochondrial properties in bivalve species with either a DUI (*Mytilus edulis* and *Ruditapes philippinarum*) or a SMI (*Mya arenaria* and *Mercenaria mercenaria*) system of mitochondrial transmission. We specifically tested: (i) whether DUI paternal mitochondria might have a constitutively higher membrane potential than the SMI ones; (ii) whether oocyte presence might induce a sudden increase in Δ*ψ*m just prior to fertilization in DUI sperm. Our results, which confirm that sperm Δ*ψ*m increases following egg detection only in DUI species, represent a key-step towards a better understanding of the mechanisms underpinning the unique ability of DUI paternal mitochondria to escape elimination and be transmitted to future generations.

## Material and methods

Adult bivalve specimens were obtained from a local fish market (Montreal, Quebec, Canada) during their spawning period, between June and August 2021. The DUI species examined were *Mytilus edulis* (Linnaeus, 1758) from Kensington (Canada) and *Ruditapes philippinarum* (Adam and Reeve, 1850) from Vancouver (Canada), representing independent origins of DUI [38]. The two SMI species were *Mercenaria mercenaria* (Linnaeus, 1758) and *Mya arenaria* (Linnaeus, 1758), both from Barnstable (USA).

Individuals were dissected on ice, gonads were excised, and sex was determined by inspecting gonadial smear under microscope. Gonads were then placed in a petri dish containing artificial sea water (ASW, 3% Instant Ocean salt mix) and mature gametes were stripped by either letting actively swim out for 5 min (sperm), or passively flow out of the gonad for 30 minutes (eggs) [33–35]. For each species, sperm were extracted from five different males and each sperm constituted a biological replicate (*M. edulis n*=1678, 1959; *R. philippinarum n*=895, 642; *M. mercenaria n*=687, 559; *M. arenaria n*=1988, 1487, control and treatment respectively).

The solution of egg-derived compounds was achieved through homogenization of mature eggs with a Polytron PT1200E (Kinematica AG, Switzerland) in ASW, (four rounds of 10s homogenization intercalated by 30s of cooling on ice). The cellular contents were then separated from bigger debris by centrifugation (10,000 RPM for 1 minute) at 4°C. The supernatant was stored at -80°C prior to experiment. For each male, motile sperm were divided into two aliquots: a control group without egg-chemoattractants (350 µL of sperm-solution, 150 µL ASW) and a group infused with the species-specific egg-derived solution (350 µL of sperm-solution, 125 µL ASW, 25 µL egg-derived solution). Sperm mitochondria were then stained with both MitoSpy™ Green FM and MitoSpy™ Red CMXRos probes (BioLegend, San Diego, CA), which respectively localize to the mitochondria in a way that is either independent (MitoSpy™ Green FM) or dependent (MitoSpy™ Red CMXRos) from the Δ*ψ*m. The final concentration for each dye was 400 nM [37]. Samples were then left to incubate at RT for 30 minutes in the dark and finally washed to remove the excess staining agent. The washing cycles consisted in removing the supernatant through centrifugation (10,000 RPM, 1 minute) followed by resuspension in 500 µL ASW, up to four times.

Cellular imagery was obtained at 60x magnification on a dedicated EVOS™ M5000 Imaging System (Thermo Fischer Scientific, USA). FITC and TRITC filters were respectively used to detect MitoSpy™ Green FM (excitation 490 nm, emission 516 nm) and MitoSpy™ Red CMXRos (excitation 577 nm, emission 598 nm). All parameters were kept constant across all captures. For each male, *n=5* separate pictures containing multiple sperm were taken for the control and another *n=5* pictures for the chemoattractant-treated sperm. Images were analysed through the Fiji distribution of Image J software [39]. Fluorescence intensity was obtained as mean grey value of the area corresponding to each sperm mitochondria, determined by automatic thresholding with the Otsu’s method (*M. Edulis, R. philippinarum, M. arenaria*). In the case of *M. mercenaria*, threshold values were manually optimized and then tested to exclude any statistical difference between treatments [40]. The area size was set up at 50-500 pixels^2^. Fluorescence intensities were corrected for the background intensity [41].

Data analysis was conducted on the software R [42]. A linear mixed model was implemented for each parameter separately, considering the factors species (*‘species’*) or inheritance (*‘inheritance’*), and presence/absence of eggs (*‘treatment’*) as categorical fixed effects, plus their respective interaction (*‘species:treatment’* or *‘inheritance:treatment’*). Factors species (*‘species’*), individual ID (*‘subject’*) and picture ID (*‘picture’*, nested within *‘subject’*) were also included as random effects. Significance was determined through a Type III ANOVA, followed by *post hoc* multi comparison with multiple testing adjustment. Statistical significance was set at *p* ≤ 0.05. Detailed summaries are provided in supplementary tables s1, s2.

## Results and Discussion

The effect of egg compounds detection on sperm mitochondrial membrane potential (Δ*ψ*m) was tested at the level of species and inheritance group (figure 1, tables s1, s2). Our analyses revealed an interaction effect between the presence/absence of egg compounds (factor *‘treatment’*) and the species tested (*F*(3,173.65)=9.6, *p*=6.665e-06***) (figure 1A). A significant increase in sperm mitochondria fluorescence intensity after treatment was revealed for the DUI species *R. philippinarum* (*p*<0.0001***). Although not significant, an increasing trend was also observed for *M. edulis*, whereas no changes were detected in both SMI species (*M. mercenaria p*=0.2952, *M. arenaria p*=0.1974) (figure 1A; table s2). In line with these results, a significant relationship between the type of mitochondrial inheritance (DUI or SMI) and the response of mitochondria to the presence of egg compounds was also detected (*F*(1,175.49)=16.4, *p*=7.66e-05***) (figure 1B). Following a multicomparison by inheritance, only the parameter DUI showed a significant fluorescence increase between control and treatment (*p*=0.0001***), whereas a non-significant opposite trend was detected for the parameter SMI (*p*=0.1071) (figure 1B; table s2). Overall, our results suggest that, along with bioenergetic changes, the modulation of Δ*ψ*m in relation to oocyte presence/detection might represent a specific property of DUI paternal mitochondria.

**Figure 1.**
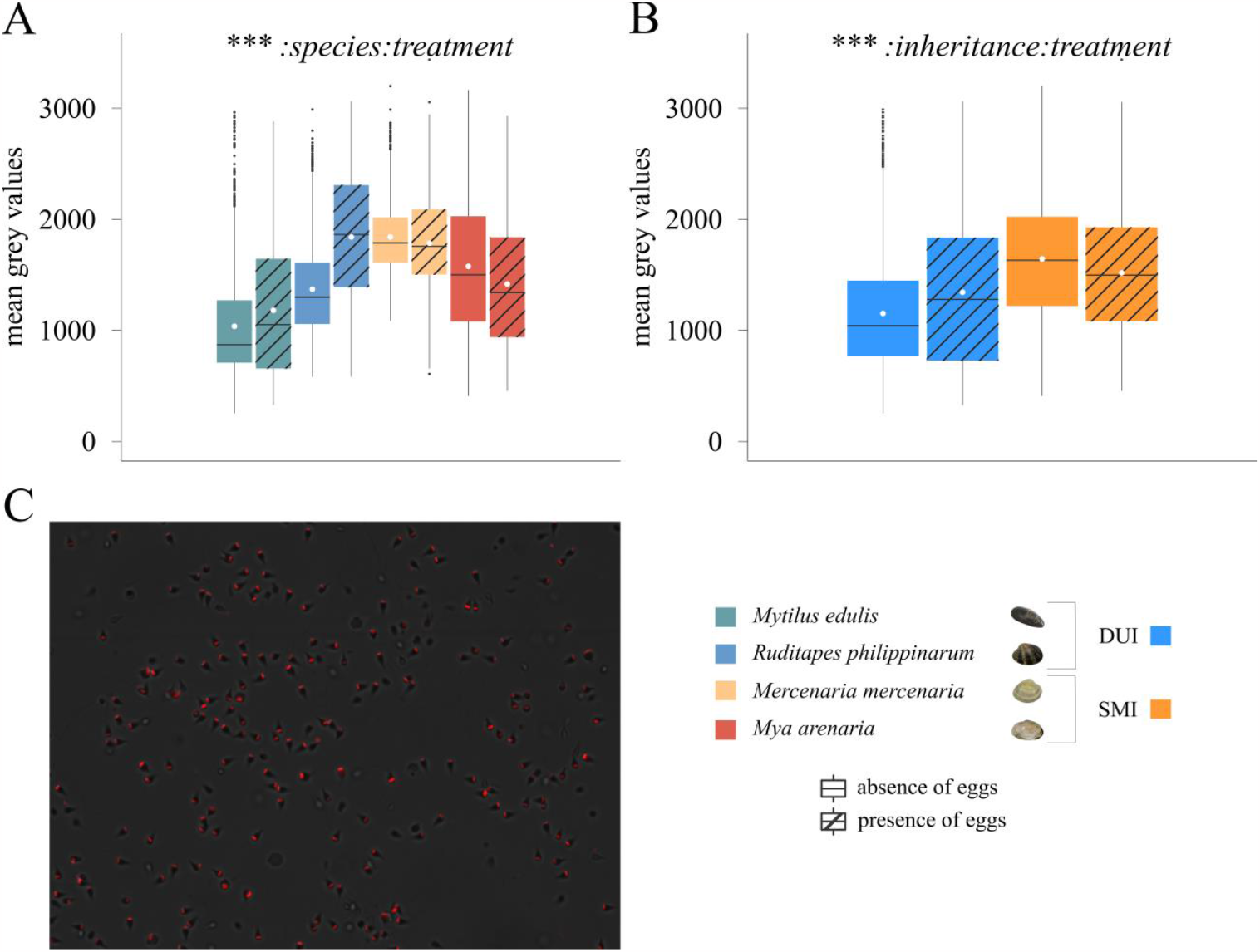
Mitochondrial membrane potential of bivalve sperm. The mitochondrial membrane potential is detected fluorometrically through the use of MitoSpy™ Red CMXRos probe and its intensity is reported as mean grey values. **(A)** Mitochondrial membrane potential across the four species analysed *M. edulis, R. philippinarum, M. mercenaria* and *M. arenaria*, in absence and presence of egg chemoattractants; **(B)** Mitochondrial membrane potential across DUI and SMI species, in absence and presence of egg chemoattractants. **(C)** Representative image of sperm mitochondria fluorescence associated with membrane potential. Statistical analyses: (A) linear mixed model with fixed effects of *‘species’* and *‘treatment’*, plus their interaction; (B) linear mixed model with fixed effects of *‘inheritance’* and *‘treatment’*, plus their interaction. Significance was determined by means of a type III ANOVA. \**p* ≤ 0.05; \*\**p* ≤ 0.01; \*\*\**p* ≤ 0.001. A detailed summary is reported in supplementary tables s1, s2.

Sperm-egg interactions through chemical signalling have been revealed to be a complex communication system with direct consequences on sperm direction, speed and even genetic selection [43–48]. However, little is known about the role played by mitochondrial and cellular bioenergetics in gamete interactions and external fertilization. Our results tend to this goal and provide evidence that egg-detection can trigger a physiological response in sperm at the subcellular level. The ability to modulate sperm Δ*ψ*m in the presence of chemical cues appears to be specific to some DUI species where paternal mitochondria are uniquely selected and transmitted to male offspring. It was hypothesized that DUI sperm mitochondria might benefit from a constitutively high Δ*ψ*m, but our results did not reveal any significant difference in Δ*ψ*m in absence of oocytes between DUI and SMI species (table s2). The dissimilarity between the two groups seems to reside in the ability to elevate Δ*ψ*m according to an external triggering factor, in this case, egg proximity (figure 1). This physiological property is in accordance with the previously acquired evidence that DUI sperm mitochondria express a male-specific phenotype able to sustain a high Δ*ψ*m. This phenotype is characterized by low respiratory rates due to a limitation exerted by cytochrome *c* oxidase and ATP-synthase complexes [32,33,35]. Together with specific mitochondrial properties, one way by which mitochondria can potentially preserve Δ*ψ*m is by reversing ATP-synthase activity at the expenses of ATP fuelled by glycolysis [49,50], or through proton release by lactate oxidation within the mitochondrial intermembrane space (see “lactate shuttle”) [51]. Although the involvement of these mechanisms in DUI Δ*ψ*m modulation is purely speculative at this point, previous work suggested that DUI sperm increasingly rely on fermentation following egg detection, with no detectable change in swimming performance [34,52]. Moreover, the low respiration rates of DUI paternal mitochondria together with the generally slow speed of sperm carrying them [34,53] suggest a mitochondrial phenotype tailored for functions that go beyond intense energy production. It is therefore possible (although yet to be proven) that this metabolic switch might provide the supply of glycolytic end products (ATP and/or lactate) needed by sperm mitochondria to modulate Δψm just before fertilization.

Our results provide new evidence that the sex-specific divergence of DUI mitogenomes underpins variation in bioenergetic properties and support the possible link between mitochondrial physiology and organelle transmission in DUI species. It is possible, yet unproven, that a sudden increase in sperm mitochondria Δ*ψ*m when in proximity with oocytes might underpin their ability to be transmitted to the offspring in a sex-specific fashion. This, through both preferential selection [3,54] and active transport into the primordial germ cells by Δ*ψ*m-dependent trafficking mechanisms [36]. The two DUI species tested are phylogenetically distant and were suggested to represent separate origins of the DUI system [38]. This reinforces the potential generalization of the results. However, further research involving a greater number of species is certainly needed.

The results presented here suggest that sperm Δ*ψ*m modulation might be involved in DUI, with potential implications in paternal mitochondria transmission. To further explore this hypothesis, we might be able to impair sperm Δ*ψ*m prior to fertilization and explore the possibility that the usual DUI aggregation pattern could be disrupted in male embryos [55–57]. We may also be able to observe a bias towards females in the offspring sex ratio if mitochondrial selection is indeed a trait involved in sex-determination in DUI species.

## Funding

This project has received funding from the Natural Sciences and Engineering Research Council of Canada (NSERC) [RGPIN-2019-04076] to S. Br. S. Br. holds the Canada Research Chair (Tier 2) in Mitochondrial Evolutionary Biology. During part of the writing/reviewing, S. Be. was supported by the European Union’s Horizon 2020 research and innovation program under the Marie Skłodowska-Curie grant agreement No 101030803.

## Notes

*Data accessibility statement:* Supportive information is provided as online supplementary tables. All data and scripts used in this study are available on figshare online repository: doi.org/10.6084/m9.figshare.24459604.v1.

### Competing Interest Statement

The authors have declared no competing interest.

